# Processing of novel food reveal payoff and rank-biased social learning in a wild primate

**DOI:** 10.1101/2020.09.25.313437

**Authors:** C. Canteloup, M.B. Cera, B.J. Barrett, E. van de Waal

## Abstract

Social learning – learning from others – is the basis for behavioural traditions. Different social learning strategies (SLS), where individuals biasedly learn behaviours based on their content or who demonstrates them, may increase an individual’s fitness and generate behavioural traditions. While SLS have been mostly studied in isolation, their interaction and the interplay between individual and social learning is less understood. We performed a field-based open diffusion experiment in a wild primate. We provided two groups of vervet monkeys with a novel food, unshelled peanuts, and documented how three different peanut opening techniques spread within the groups. We analysed data using hierarchical Bayesian dynamic learning models that explore the integration of multiple SLS with individual learning. We i) report evidence of social learning compared to strictly individual learning, ii) show that vervets preferentially socially learn the technique that yields the highest observed payoff and iii) also bias attention toward individuals of higher rank. This shows that behavioural preferences can arise when individuals integrate social information about the efficiency of a behaviour alongside cues related to the rank of a demonstrator. When these preferences converge to the same behaviour in a group, they may result in stable behavioural traditions.

## Introduction

Organisms may learn about their environment and acquire new skills individually, by trial and error, or socially, by directly observing and/or copying others’ behaviour [1]. Social learning has been considered adaptive since organisms may gain fitness benefits when learning from others, especially in a changing environment or when individual learning is costly [2]. However, social learning might also be maladaptive when social information is outdated [3, 4]. Socially transmitted behaviours resulting in animal cultural traditions have been described in diverse taxa and domains such as songbird vocal dialects [5], tool use in crows [6], hunting techniques in whales [7], social conventions, extractive foraging techniques and manipulation behaviours in numerous primates [8–11].

Social learning is often not random, and animals can use social learning strategies to acquire behaviour. Social learning strategies are rules of thumb that permit an individual to acquire behaviour rapidly or cheaply without evaluating the entire population or understanding the functional significance of a behaviour. These strategies can also structure within- and between-group variation in behavioural traditions. Social learning biases direct an individual’s attention towards a behaviour based on its content, relative frequency in a population, or its association with a particular type of demonstrator. These biases are typically, but not always, assumed to be shortcuts to acquiring adaptive behaviour and are theoretically well explored [3, 12, 13]. Content-biases focus on characteristics of the observed behaviour such as a bias towards more successful behaviours, which yield better payoffs.

Much theoretical work has explored the evolution of payoff-biased learning [14, 15], which is useful when payoffs are stochastic or adaptive behaviour is rare. Empirical work in humans show payoff-bias is used concurrently with other social learning strategies [16]. Researchers have recently expanded their investigation of payoff-biased learning to wild animals. Wild capuchins adopted the most successful and efficient technique when processing novel fruits [17]. Great tits do not use payoff-biased learning but acquire a high payoff foraging technique through a combination of positive frequency-dependent learning and individual learning that is highly sensitive to personal experiences in payoffs [18]. Male vervet monkeys, but not females, more often used the technique displayed by male demonstrators when a difference in payoff was introduced with males receiving more food than females [19]. The prevalence of payoff-biased learning is likely underestimated in wild populations particularly if adaptive behaviour is common, easy to innovate, or if there is a single optimal solution [17].

Context biases focus on other cues such as their frequency in a population (e.g. preferentially use the most common behaviour or the one exhibited by the majority of individuals: conformity bias) or particular traits of models (e.g. preferentially use the behaviour displayed by familiar, prestigious, related, or similarly aged individuals: [20]). Such biases have been reported in various animal species. “Moss sponging” by chimpanzees [21] and “sponging” by dolphins [22] were both transmitted within matrilines; “shelling”, another foraging technique, spread non-vertically among associated dolphins [23]. Younger female guppies socially learned the mate choice of older females [24]. Capuchins and chimpanzees used the foraging method displayed by older individuals [17, 25]. In female- philopatric vervet monkeys, females were preferentially chosen over males as demonstrators [26], but demonstrators were all of high-rank. The techniques used by higher-ranked individuals were preferentially chosen by chimpanzees [27] and vervet monkeys [28] when researchers simultaneously tested for several biases. Animals also conform to the most frequently observed behaviour: in vervet monkeys, dispersing males conformed to the local foraging norm, abandoning their natal preference [29]. Great tits and chimpanzees disproportionately adopted the most frequent local foraging technique [30, 31 but see 32]. Nine-spined sticklebacks preferentially chose the feeder favoured by the majority of individuals, even when this feeder was less rewarding than an alternative [33]. Fruit flies chose their mating partner according to the preference of the majority of copulating conspecifics [34]. All these findings suggest that different contextual characteristics strongly influence social learning patterns. However, relatively little research has investigated how organisms jointly integrate multiple social learning strategies with individual experience. This integration is important as combining social and individual learning is more adaptive than solely relying on one or the other [35].

Open diffusion experiments with experimentally trained demonstrators in the wild have been well utilized [18, 29, 30], and show what social learning strategies animals are capable of in a well-defined experimental context. However, open diffusion experiments without experimentally trained demonstrators may better show how traditions endogenously arise in the wild in more realistic social settings; yet they are rare [17, 28].

Recently developed dynamic statistical models including Network Based Diffusion Analysis (hereafter NBDA: [36, 37] and Experience-weighted Attraction models (hereafter EWA: [15, 17, 38]) are powerful statistical modelling approaches to identify evidence for social learning and the pathways of social transmission. NBDA is useful to identify evidence for social learning and the typical pathways of social transmission, but it only analyses the initial spread of a new behaviour, i.e. the origin of a cultural tradition. It cannot thus evaluate what social learning strategies individuals use to inform behavioural choice once they have learned multiple options [37]. Nor can NBDA examine how individuals uniquely combine individual and social information – the combination of which makes social learning adaptive. EWA models analyse the entire unique behavioural sequence chosen by each individual and investigate how this is influenced over time by both personal experience and observation of others, i.e. the origin and maintenance of a cultural tradition [15, 17]. EWA models permit extant theoretical models of social learning strategies to be turned into statistical models to examine which strategy, combination of strategies, or individual learning alone best predict an individual’s behaviour. These predictions are conditional upon an individual’s unique personal experiences and opportunities for observing social information, thus linking individual variation in behaviour to population-level cultural dynamics.

This study aims to determine which content and/or context biased social learning strategies are responsible for the transmission of novel food processing techniques - peanut shelling - in wild vervet monkeys (*Chlorocebus pygerythrus*). We tested for payoff-biased learning as content-biased strategy and for frequency-dependent and model-based learning (sex, kin and rank) as context-biased strategies. Here, we offered a novel food – unshelled peanuts – to two groups of wild vervet monkeys: Noha (hereafter ‘NH’) and Kubu (hereafter ‘KB’) during four months of field experiments. We recorded the exact time of each peanut consumption event and the identity of observers at each time to construct a dynamic observation network. We described the three different peanut extraction techniques used by monkeys: crack with the hand (hereafter ‘CH’; Movie S1: https://youtu.be/BkiTJyxp3gQ); crack with the mouth from the side of the peanut (hereafter ‘CMS’; Movie S2: https://youtu.be/3JOAH4khnjk) and crack with the mouth from the top of the peanut (hereafter ‘CMT’; Movie S3:: https://youtu.be/RaFoQpBadG8). We then analysed data from this experiment with EWA models. We predict, in accordance with another study focusing on context biases [28], that higher-ranked vervet monkeys are more influential demonstrators than others. We hypothesize that other context biases such as frequency-dependence or content biases such as payoff-bias (i.e. the most successful technique), would also be possible. The major interest of this study is to assess whether vervets rely purely on individual learning, or integrate individual learning with social learning strategies, by fitting multiple hierarchical EWA models to data collected in an open diffusion field experiment with no trained demonstrator.

## Results

### Effects of group, rank, sex and age on success, manipulation and observation

In NH, 25 individuals (9 adults; 12 juveniles; 4 infants) succeeded in opening 2104 peanuts and attempted to open a further 1401 peanuts. In KB, 9 individuals (5 juveniles; 4 infants) succeeded in opening 45 peanuts and 10 individuals (6 juveniles; 4 infants) attempted to open 102 peanuts (Table S1). We found an effect on rank and sex both on the rate of peanut opening successes and on the rate of manipulation. Higher rankers succeeded significantly more to extract peanuts (GLMs: β estimate=-2.67; standard error (SE) =0.67; t value=-4.00; P=0.0004) and manipulated more peanuts than lower rankers (β=-2.22; SE =0.65; t value=- 3.56; P=0.002). Males succeeded significantly more to extract peanuts (18/24 males; 16/29 females; β=1.12; SE =0.32; t value=3.56; P=0.001) and manipulated significantly more peanuts than females (19/24 males; 16/29 females; β=0.93; SE =0.32; t value=2.95; P=0.007).

The latency of first success ranged from 2.03 minutes to 448.1 minutes in NH (mean=218.35 min; SE of the mean=25.29 min) and from 78.95 min to 393.6 min in KB (mean=223.33 min; SE of the mean=38.85 min; Table S1). The innovator, in terms of opening peanut, in NH was a low-ranked, newly immigrated adult male at the first exposure to peanuts (Avo) while, in KB, it was an infant male (Aar) at the third exposure. We found a significant effect of age on latency of first peanut opening success. Juveniles had a lower latency compared to infants (GLMs: β=-0.57; SE =0.25; t value=-2.34; P=0.05), but we found no significant difference between infants and adults (β=0.13; SE =0.31; t value=0.41; P=0.91) nor between juveniles and adults (β=-0.45; SE =0.25; t value=-1.82; P=0.16). We found no significant difference between groups (β=-0.16; SE =0.24; t value=-0.67; P=0.51), between males and females (β=0.12; SE =0.21; t value=0.57; P=0.58) and no significant effect of rank (β=-0.11; SE =0.42; t value=-0.26; P=0.80) on latency of first success.

We found no clear difference in the rate of observation of successes between groups (GLMs: β=2.79; SE =1.66; t value=1.68; P=0.10), between high rankers and low rankers (β=- 1.64; SE =0.86; t value=-1.91; P=0.07), between adults and infants (β=0.64; SE =0.59; t value=1.08; P=0.29), between adults and juveniles (β=-0.007; SE =0.42; t value=-0.017; P=0.99) nor between males and females (β=0.81; SE =0.44; t value=1.85; P=0.08). We found limited evidence for any difference on the rate of observation of manipulations between groups (β=2.42; SE =1.36; t value=1.78; P=0.09), between high and low rankers (β=-1.56; SE =0.83; t value=-1.89; P=0.07), between adults and infants (β=0.57; SE =0.58; t value=0.98; P=0.33), between adults and juveniles (β=-0.01; SE =0.41; t value=-0.03; P=0.98) and between males and females (β=0.79; SE =0.42; t value=1.88; P=0.07).

We found effects of rank, age and sex on the rate of being observed when manipulating and successfully opening peanuts. High-rankers were observed significantly more than low-rankers when succeeding (GLMs: β=-3.61; SE =0.65; t value=-5.57; P<0.001) and manipulating (β=-3.31; SE =0.65; t value=-5.08; P<0.001). Adults were observed significantly more than juveniles when succeeding (β=-0.91; SE =0.29; t value=-3.12; P=0.004) and manipulating (β=-0.73; SE =0.29; t value=-2.56; P=0.02). Males were observed significantly more than females when successfully opening (β=1.15; SE =0.27; t value=4.20; P<0.001) and manipulating peanuts (β=1.02; SE =0.28; t value=3.63; P=0.001).

### EWA model interpretations

For most individuals, and at the population level, we found that CMS was the most successful technique, and CMS became the most common technique in the population. CMS was successful on 46.8% occasions in NH and 24.8% in KB over all manipulations. This was followed by CMT (8.1% in NH, 5.4% success in KB) and CH (5.1% in NH, 0.7% in KB). From WAIC values and inspecting model predictions against raw data, we found that the global model best predicts our data compared to any model representing a single social learning strategy (Table 2) and successfully predicts the frequency of each technique used at the population-level (Fig. 1) and across all individuals (Fig. 2; Fig. S1). Although the global model has the lowest WAIC score and predicts data best across all individuals in the population, there is more to learn from comparing predictions and parameter estimates from all models as it helps us better understand the importance of different cues and their corresponding learning strategies. Model predictions show that the highest-payoff behaviour became the most common for nearly all individuals in the population (Fig. 2; Fig. 3 and Fig. S1). The parameters’ magnitudes and certainties in the global model, as well as WAIC score and parameter estimates in individual models, suggest that payoff-bias, followed by rank-bias, were the two singular most important learning strategies (Table 2; Table 3).

**Figure 1.**
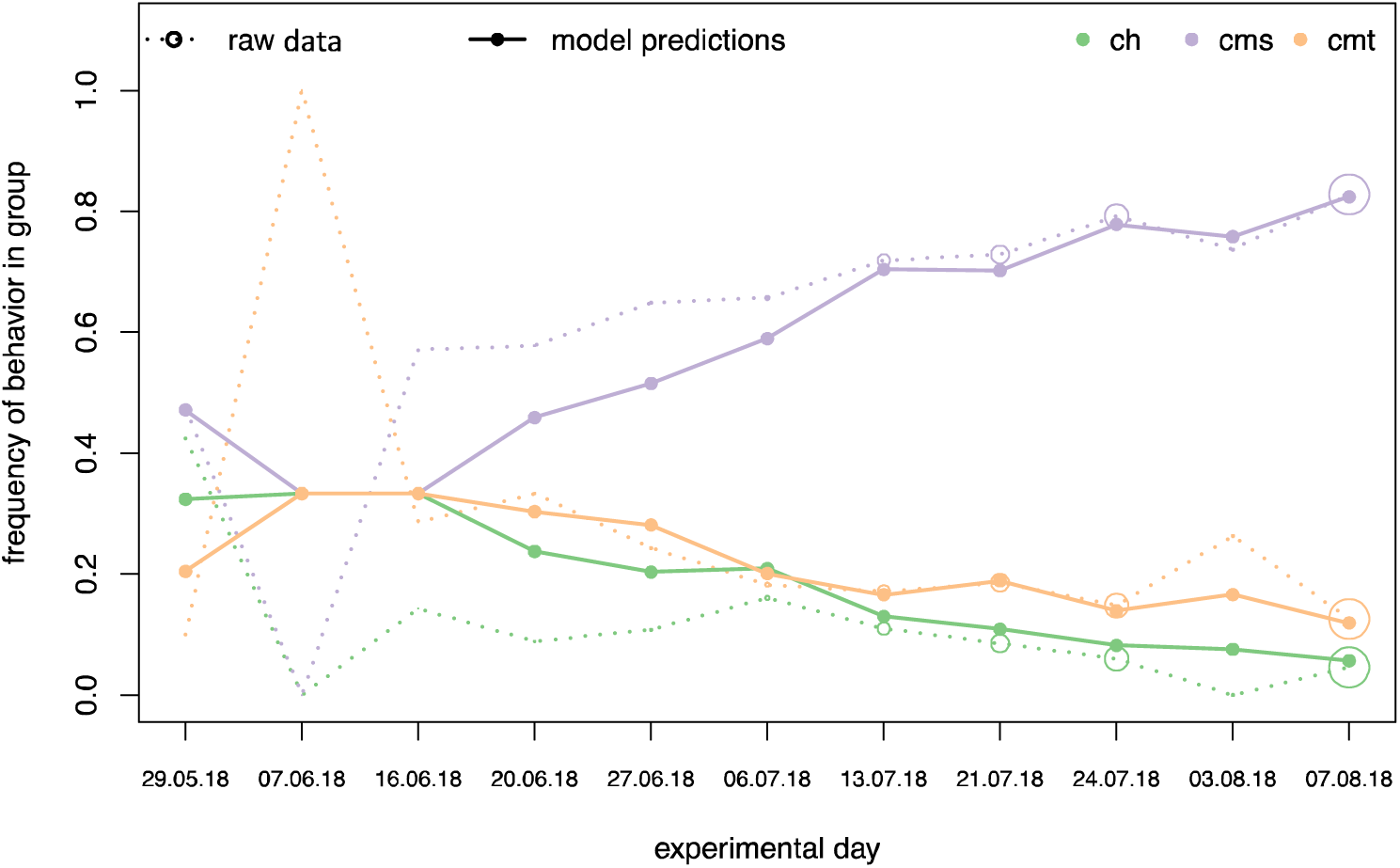
Daily mean group probability of observed processing techniques over experimental days for NH. Filled lines and points are posterior mean predictions of the global model averaged across all individuals in NH in an experimental day. Dashed lines and empty points are estimates from raw data averaged across all individuals in NH in an experimental day. Diameter of empty points of raw data scales with daily sample size.

**Figure 2.**
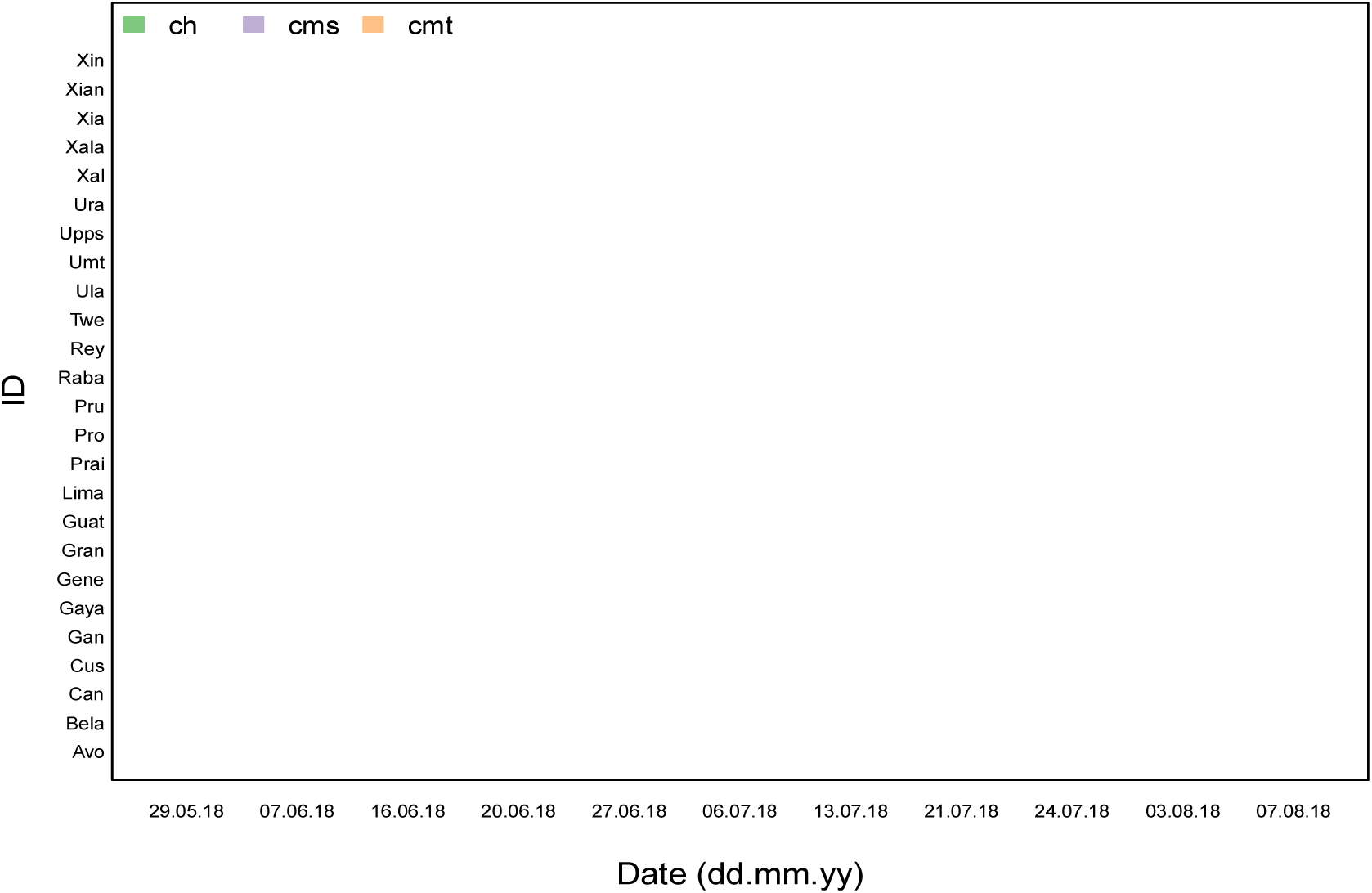
Visualization of posterior estimates from the global model of daily mean probability of processing techniques across all foraging individuals in NH. Point diameter scales with probability, with many individuals approaching fixity of CMS on the final day.

**Figure 3.**
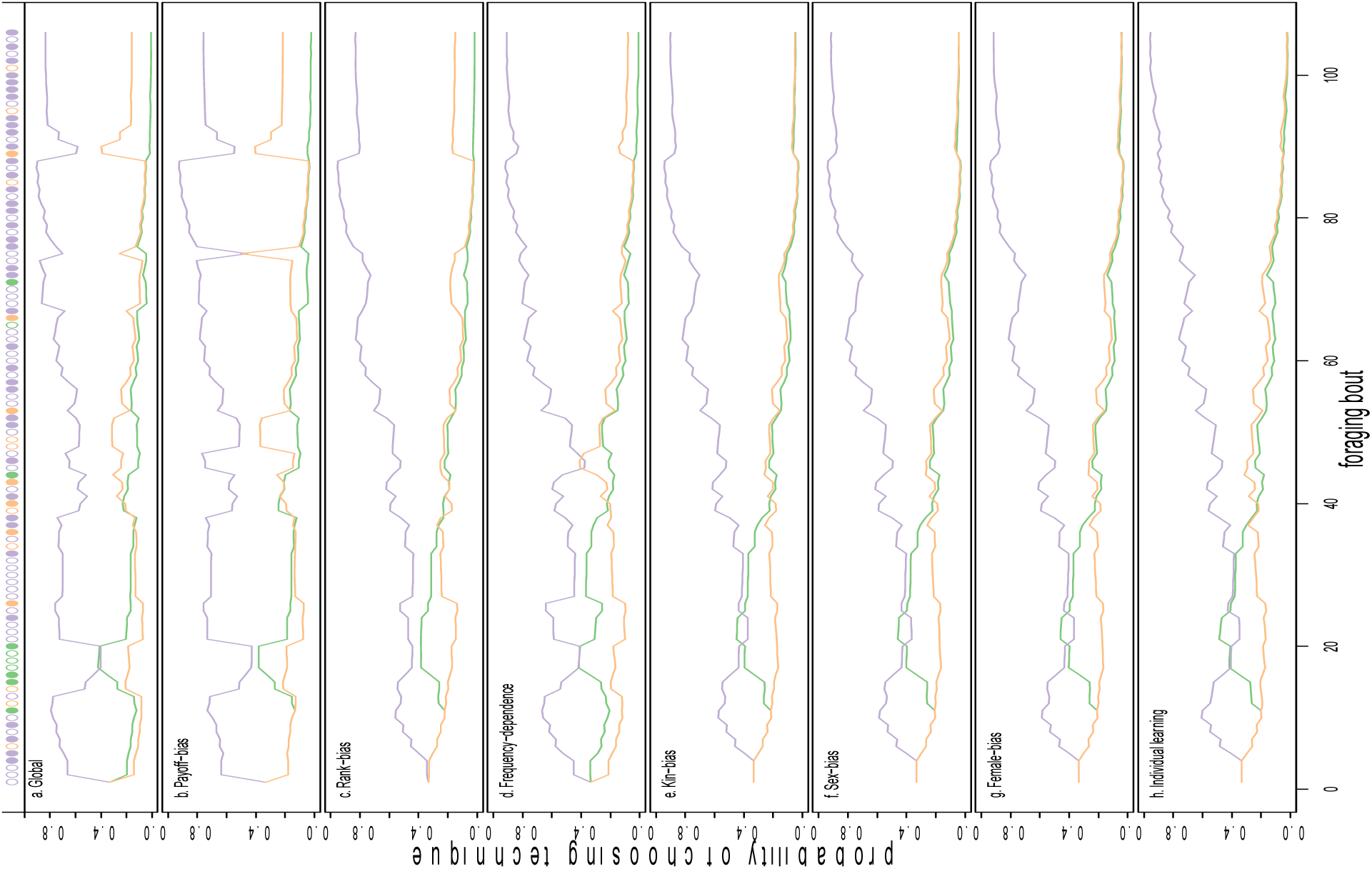
Model predictions of the probability of using a technique across foraging bouts from the seven models for the behavioural time series of individual ‘Gran’ in NH group. Colours correspond to the three processing techniques (purple: ‘CMS’; orange: ‘CMT’; green: ‘CH’). . Lines show posterior mean predictions of the probability of choosing a behaviour at each time step. Lines are like three-sided dice whose odds change as a function of observed social and experienced personal information over each foraging bout. The “roll” of each die estimates the processing technique observed by ‘Gran’ at each foraging bout, plotted along the top row. Filled circles are successes, empty circles are failures The shaded interval is a 89% High Posterior Density Intervals (HPDI). 89% HPDI correspond to the range containing 89% of probable values.

**Table 1.** Individual information and social cues used in the EWA models. Models 2-8 also incorporate individual experience from model 1. Information observed at timestep t for models 2-8 was the mean of each cue, κ evaluated in the 20 minutes prior to timestep t.

| Model | Social and Individual Cues |
| --- | --- |
| 1) Individual learning | $\pi_{ij,t} = 1$ if successful, 0 if not |
| 2) Frequency-dependence | $N_{ij,t}$ = count of times $j$ observed demonstrators perform behavior<br>$i$ |
| 3) Female-bias | $\kappa_{k,ijt} = 1$ if adult female, 0 otherwise |
| 4) Matrilineal kin-bias | $\kappa_{k,ijt}$ = coefficient of relatedness, $r$, between observer $j$ and<br>demonstrator |
| 5) Payoff-biased<br>learning | $\kappa_{k,ijt}$ = mean observed payoff, $\overline{\pi_{ij,t}}$, of behavior $i$ observed by $j$ |
| 6) Rank-bias | $\kappa_{k,ijt}$ = rank of demonstrator determined by I&SI method |
| 7) Sex-bias | $\kappa_{k,ijt} = 1$ if same sex as observe, 0 if different |
| 8) Global model | all cues in 1-7 |

**Table 2.** Widely applicable information criteria estimates for all evaluated models with a 20 minute window of social information. Standard errors, difference in WAIC and model weight are included.

| **Model** | **WAIC** | **SE** | **dWAIC** | **weight** |
| --- | --- | --- | --- | --- |
| Global Model | 4707.71 | 91.09 | 0.00 | 1.00 |
| Payoff-bias | 4756.90 | 91.12 | 49.18 | 0.00 |
| Rank-bias | 4780.21 | 90.55 | 72.50 | 0.00 |
| Frequency-dependence | 4796.59 | 90.92 | 88.88 | 0.00 |
| Matrilineal Kin Bias | 4856.82 | 92.46 | 149.11 | 0.00 |
| Same-sex-bias | 4874.49 | 92.68 | 166.78 | 0.00 |
| Female-bias | 4875.25 | 92.73 | 167.54 | 0.00 |
| Individual learning | 4920.83 | 92.00 | 213.12 | 0.00 |

**Table 3.**
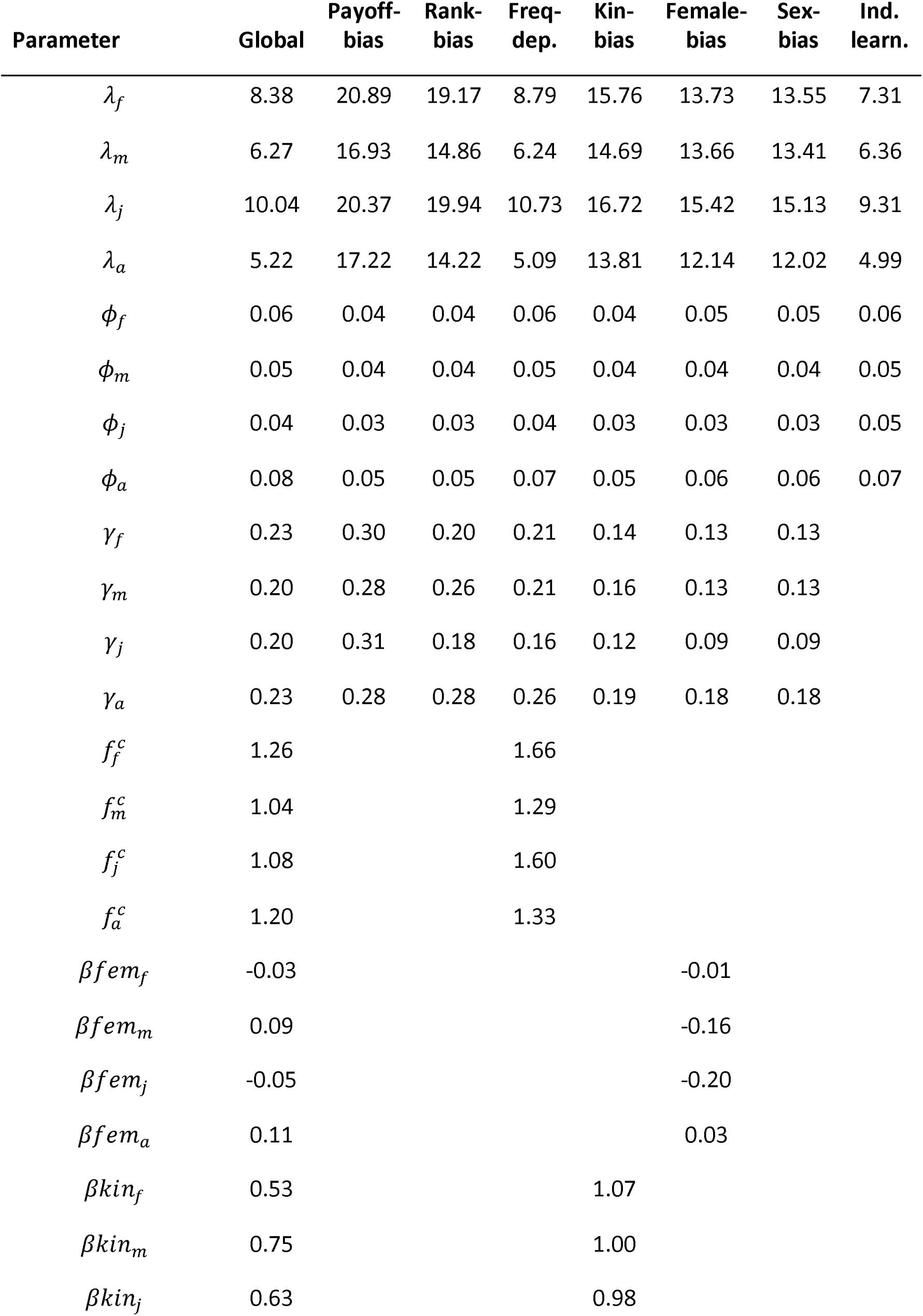
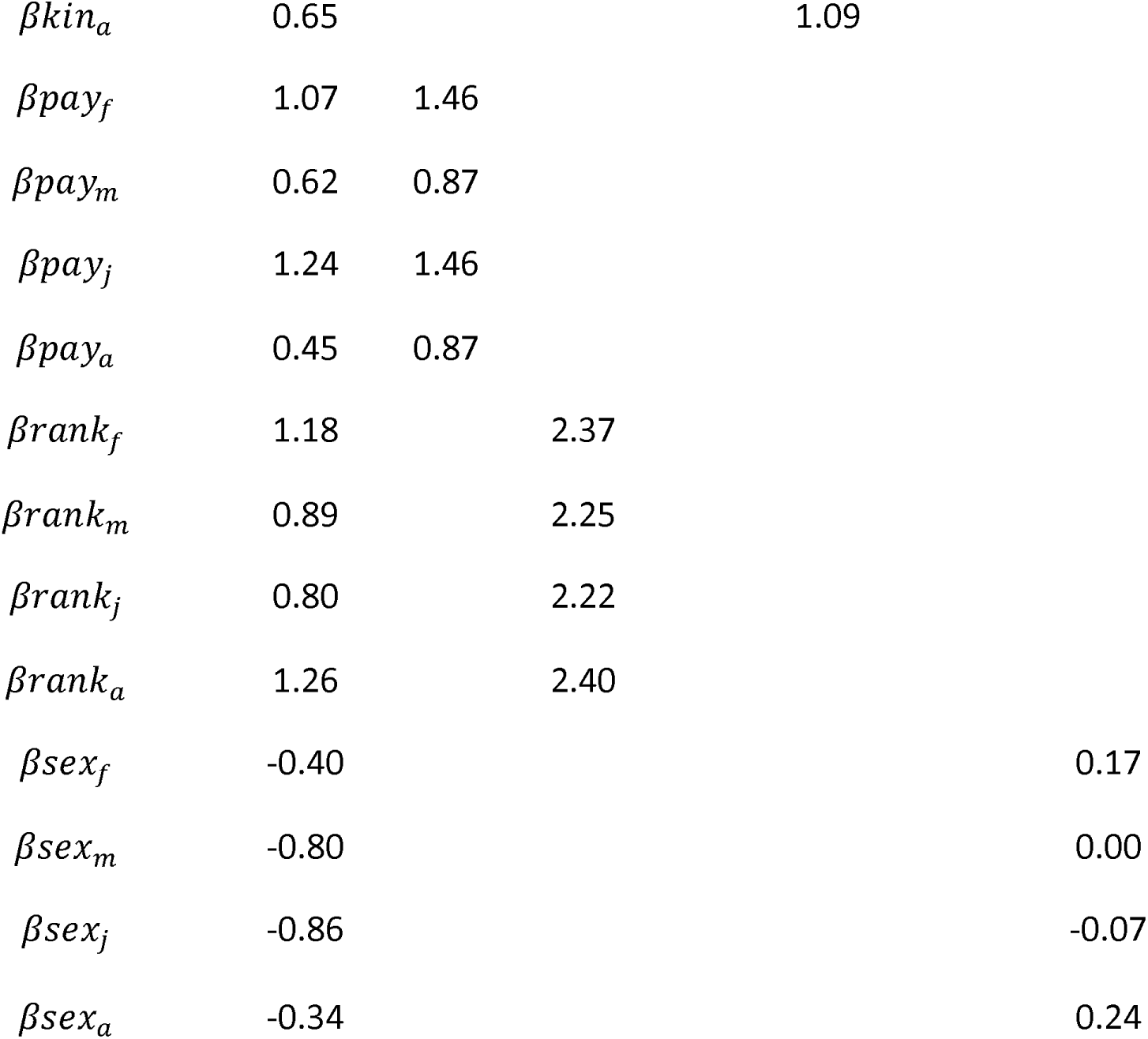
Posterior mean estimates of learning parameters for all evaluated EWA models. Subscripts correspond to male (m), female (f), adult (a), and juvenile (j). Dot plots of these parameters, including 89% HPDI for the global model, including individual and group-level parameters, are in Figures S2-S5.

Figure 3 shows predictions for the seven evaluated models for a single individual (‘Gran’ in NH). In this case, all EWA models allow the highest payoff behaviour (CMS; displayed in purple) to become the most common in the population. However, their dynamics and ability to accurately predict the behavioural sequence differ. The global model tends to show more uncertainty around any particular prediction, whereas models evaluating single learning strategies tend to be overconfident in their estimates, less accurate across the behavioural trajectory, or do not predict well for all individuals in the population. Payoff-biased and rank-biased learning models perform well, but do not adequately predict the frequency of all three behaviours across the entire time series, whereas frequency-dependent learning over-predicts the frequency of the most common, high-payoff behaviour. The individual learning model is the one having the least support from WAIC values (Table 2) and visual inspection of model predictions, strongly suggesting that individual learning alone does not explain techniques’ spread. The individual learning model’s predictions are overconfident, tend to drift around, and underestimate the frequency of CMS, particularly at the beginning of the time series. Reproducible code to plot these predictions from all models for all individuals in the population is found in the online repository given in the Method section.

We found little support that individuals preferentially used the technique displayed by females or individuals of the same sex (Table 2). Kin-biased learning might be important for a few, but not most individuals. We encourage readers to examine Figure S1, which shows global model predictions of behavioural choice for each individual vervet, as we believe it is the most informative about model fit compared to real data and is informative about individual variation. Figures S2-S11 show dot plots of parameter predictions of the global model for age category, sex, group, and individual.

### Variation between individuals and groups in parameters

We see considerable variation between the two groups and among individuals for many varying effects parameters in these models, this occurred for several reasons. In KB, fewer individuals participated in the experiment and they had fewer foraging bouts. This generated more uncertain estimates as reflected by width of the HPDIs of parameter prediction for KB compared to NH, visualized in green in Figures S2-S11. They were also less likely to observe social information, based on the 20-minute time windows we selected in this analysis. For these reasons, many of the individual-level parameter estimates of varying effects have tighter 89% HPDIs than the population mean estimates.

Posterior estimates of group-level varying effects also differ greatly for some parameters. These estimates are more precise and parameter values related to social learning are higher for NH due to the greater number of individuals who participated to the experiment. This can be visualized when looking at the shaded HPDIs around model predictions in Figure S1 for individuals in both groups of different sample sizes. It is also visualized in Figures S2-S10. Estimates of σ, the variance estimated between groups and individuals for each parameter, are extracted from the variance-covariance matrix of the global model are in Figure S11.

### Sensitivity to attraction scores (λ)

Females (Posterior Mean (λ_f_=8.38; [89% HPDI: 4.01-14.86]) are more sensitive than males (λ_m_=6.27 [3.35-10.35]) to differences in attraction scores while juveniles are more sensitive than adults (λ_j_ =10.04 [5.45-17.07]; λ_a_ =5.22 [2.64-9.03]). A higher attraction score typically means that individuals are more sensitive to personally-experienced differences of behavioural payoffs. The direction of these effects is consistent across nearly all evaluated models (Table 3). We see considerable variation in varying effects of λ at the group and individual level (Fig. S2), but much of this is due to sampling differences.

### Weight given to recent experiences (□)

Overall, we see small values for □, suggesting that individuals tend to weight past experiences more heavily than recent experiences; memory strongly influences behaviour. Adults tend to weight new experiences more heavily than juveniles (□_a_=0.08 [0.02-0.17]; □_j_=0.04 [0.01-0.08]). We see little support for sex differences (□_f_ =0.06 [0.02-0.12]; □_m_=0.05 [0.01-0.11]). The direction and magnitude of these effects are consistent across all evaluated models (Table 3). Groups and individuals (particularly in KB) tend to have larger and more uncertain estimates of □ as a wider range of parameter values can predict the observed data (Fig. S3).

### Weight given to social information (γ)

In the global model, social information influences behaviour slightly more heavily for adults than juveniles (γ_a_=0.23 [0.05-0.48]; γ_j_=0.20 [0.05-0.38]), and for females than males (γ _f_ =0.23 [0.05-0.43]; γ_m_=0.20 [0.04-0.43]). However, these patterns are not consistent across all models. This is because for each individual model, one learning bias might be particularly salient for a particular age or sex class. We can gain insight into this from comparing predictions from the global model to models representing individual learning strategies (Fig. S4). For example, the payoff-bias learning model has higher values of γ for juveniles and females (γ_j_=0.31 [0.13-0.50]; γ_a_=0.28 [0.11-0.49]; γ_f_ =0.30 [0.13-0.51]; γ_m_=0.28 [0.11-0.49]), while the rank-bias learning model has higher values for of γ for adults and males (γ_a_=0.29 [0.12-0.50]; γ_j_=0.18 [0.07-0.33]; γ_m_=0.27 [0.11-0.47]; γ _f_ =0.20 [0.08-0.37]).

### Payoff-bias

We find reliable support for payoff-biased learning, for nearly all individuals in the population (Fig. S6). Of the social learning biases, we also see the most salient differences between age and sex classes. Juveniles rely more on payoff-biased social learning than adults (βpay_j=_1.24 [-0.36-2.76]; βpay_j_=0.45 [-1.06-1.97]), while females rely more heavily on it than males (βpay_f_=1.07 [-0.49-2.63]; βpay_m_=0.62 [-0.88-2.08]). Of the EWA models representing single social learning strategies, payoff-biased learning had the lowest WAIC value (Table 2) and its predictions were most similar to the global model for most individuals in the population. Similar differences and stronger effect sizes were estimated in the payoff-biased learning model when compared to the global model (β_pay_ in Table 1).

### Rank-bias

We also see good support for rank-biased social learning for most individuals in the population, although there is more uncertainty around these predictions (Fig. S7). The global model suggests that adults are more likely to learn from high-ranking individuals than juveniles (βrank_a_=1.26 [-0.25,2.72]; βrank_j_=0.80 [-0.85,2.53]), and females are more likely to learn from high-ranking individuals than males (βrank_f_=1.18 [-0.40,2.75]; βrank_m_=0.89 [- 0.60,2.48]). These differences between age and sex parameters are much smaller in the rank-biased learning model (See βrank in Table 3). This is likely because higher ranking individuals are more successful than low ranking individuals. When including both rank and payoff in the global models, differences in reliance on rank cues, instead of payoff cues, emerge.

### Frequency-dependence

On average, the population tends to be somewhere between linear and slightly positive-frequency-dependent (See f^c^ in Table 1). We see posterior means slightly larger than 1 for most individuals (Fig. S5). However, boundary conditions (f^c^>0) often make posterior point estimates misleading; nearly half of the posterior mass for many individuals and all age and sex classes lies in a parameter space consistent with anti-conformity. The frequency-dependent learning model estimated a stronger degree of conformist transmission, but it often under-predicts the probability of an individual choosing a less common behaviour (Figure 3).

### Matrilineal kin-, Female- and Same-sex biased learning

We see some evidence for matrilineal kin-biased learning (See β_kin_ Table 3), but this might only be for some individuals in the population (i.e. Pro, Gran, Xian in Fig. S8). We see little evidence for sex-biased and female-biased transmission (Table 3; Fig. S9; Fig. S10).

## Discussion

In the present study we used a time-series model predicting each technique’s probability of being chosen to infer which social learning strategies were utilized in two groups of wild vervet monkeys when learning to process unshelled peanuts, a novel food. The best supported statistical model, and evidence from a suite of models integrating individual and social learning, suggests that vervet monkeys learned the most successful, highest payoff technique, and displayed by the higher ranked demonstrators. We found weak support for a positive frequency-dependent conformity bias defined as ‘use the most observed technique’. We also found no clear evidence that they show a bias toward the technique displayed by females, individuals of the same sex, or related individuals. Individual learning alone is very unlikely to be responsible for the techniques’ spread.

We reported high variation between groups and among individuals for many parameters in the models (Figure S11). This group-level variation is likely due to differences in data sample size between the two groups: one group (NH) with many more participants eating peanuts than the other (KB). Consequently, estimates of parameters for individuals in KB were more uncertain and often had larger posterior means. These larger point estimates for lambda, phi, gamma, and fc are likely not due to group-level differences but likely arise because they are near a boundary condition of zero, and uncertainty drives mean estimates upward. It is thus important to evaluate the full shape of posterior and be cautious before drawing conclusions from point estimates of parameter differences between groups and individual with unequal sampling. Finally, we reported effects of age, sex and rank on individual and social learning which are discussed in the remainder of the discussion.

Note that we avoided using the term “copying” across the manuscript which is strongly associated to “imitation” by some scholars [39, 40], although not by all [11, 41]. Here, we do not make any assumption and conclusion regarding the social learning mechanisms (action form copying versus non-form copying/socially mediated individual learning: [40]) implied in the acquisition of the peanuts processing techniques. We acknowledge that the three peanuts processing techniques we describe are within the Zone of Latent Solution of vervet monkeys [40] and we do not claim for form copying, i.e. that vervet monkeys faithfully copied the exact movements – the form of the behaviour - displayed by their conspecifics.

We found the strongest support for the global model suggesting that vervet monkeys use multiple social learning strategies simultaneously. Comparison of parameter effect sizes across models, model predictions, and WAIC values all suggest that payoff- and rank-biased learning guided diffusion of peanut processing techniques in both groups of wild vervet monkeys. These results do not fully align with those of a previous study on vervet monkeys at our field site [19] that can be explained by methodological differences. In Bono et al. study, payoff was operationalized as a different quantity of food provided by a two-action design – one item versus five. In our study, socially observed payoffs were the mean observed probability of a technique being successful in the previous 20 minutes. While only observations of trained demonstrators were taken into account in Bono et al. study, here observed payoffs are unique to each individual’s full observation experience. Bono et al. found that only males chose the technique used by the male model more when it received five food items compared to a female model receiving only one item [19]. We find that females and juveniles relied more on payoff-biased social learning than males and adults conversely. We believe this discrepancy in results can be explained by methodological differences between both studies. First, in Bono et al. study [19], only two models, a male and a female, that were both adults and of high rank, were available in each group, while in our study, numerous individuals of both sexes and various ages and ranks could opportunistically serve as demonstrators. Second, we tested behavioural responses of individuals towards a novel and unknown food while Bono et al. [19] tested monkeys in a two-action box experiment filled with known food. It is possible that monkeys got accustomed to boxes’ affordance across the numerous experiments of this kind they had previously experienced at the Inkawu Vervet Project. Different strategies could be at play in different environmental contexts, i.e. in a familiar foraging situation *versus* a novel one. Our results suggest thus that females - the philopatric sex – are maybe less conservative in their social learning than previously thought and that their conservatism decreases in a novel context after an environmental change. When it has been suggested that females would stick to their habits compared to males who are expected to show more behavioural flexibility [19], our findings challenge this hypothesis. Indeed, in vervet monkeys, females are the philopatric sex, meaning that they remain all their life in their social group while males disperse to new groups several times within their lifetime. Females possibly possess thus more detailed knowledge than males about the distribution of existing food resources or objects such as boxes they encounter in their territory, which could make them a valuable source of directed social learning in these common situations. However, we suggest that when facing an unknown foraging situation, females can show flexibility, decrease their conservatism and rely more than males on the social information available.

Other recent studies used a similar multilevel dynamic learning model in great tits [18] and in capuchins [17]. Great tits have been found to switch to an alternative higher payoff variant when given a choice between a low and a high payoff option [18]. As we found in vervet monkeys, young birds were more likely to use social information than older birds [18]. Barrett et al. [17] offered *Sterculia apetala* fruits to a naïve group of wild capuchins and followed the spread of the processing techniques. They found that, of multiple biases, payoff- biased social learning was the one responsible for the spread of the most successful techniques [17], and that age was a strong predictor in the heterogeneity of learning among individuals. We can hypothesize that this maintained preference for CMS technique could be at the origin of cultural transmission of peanut processing techniques if this food was made available on the long term in vervet monkeys’ environment and if other groups developed a preference for another technique.

Here, we also report a rank bias in agreement with the ‘use preferentially the behaviour displayed by higher-ranked individuals’ strategy found in chimpanzees [27] and vervet monkeys [28]. In this latter study, we used another modelling approach – NBDA – to identify the typical pathways of transmission of boxes opening techniques in an open diffusion experiment. We tested whether the diffusion followed specific social networks representing different pathways of learning such as learning from females, older individuals, related individuals or higher rankers. We discovered evidence of a transmission bias favouring learning from higher-ranked individuals, with no evidence for age, sex or kin bias [28]. Present results are thus in accordance with these previous findings. In this previous study, we could not test for frequency-dependent learning and payoff-biased learning, nor for the interplay between the use of social and individual information, but this is precisely what we assessed here. This means that higher rankers are not only considered as models for the first learning event but that this bias is maintained across time.

While higher rankers were manipulating more boxes and were more successful, they were not observed more than low rankers in our previous study [28]. Instead, observations of higher-ranked individuals had a greater effect on observers’ behaviour than observations of lower-ranked individuals. Here, we found that higher rankers manipulated and successfully processed more peanuts, probably because they can monopolize access to food. We did not find any difference in observation rate between individuals but, we reported that higher rankers, adults and males were observed more than lower rankers, juveniles and females. This difference might be due to experimental novelty arousing a different attentional pattern. Indeed, in a box experiment, vervet monkeys showed a selective attention to female models [26] when all models were of high rank. This is in accordance with literature on chimpanzees and capuchins in which older and more proficient nut crackers are more observed than others [42, 43].

We tested whether individuals showed a preference for the most frequent or the least frequent behaviour, or whether they showed no frequency bias. We found little evidence for any frequency-dependent learning, our population being slightly positive frequency-dependent. A previous study on vervet monkeys reported that migrant males conformed to their new group norm by abandoning their natal feeding preference, thus following a functional definition of conformity [29]. The authors’ claim of conformity [29] has been considered a premature conclusion regarding the mechanism of conformity, as the authors did not have the observational data of who the males attended to before switching and could not prove that the males observed the foraging choice of a majority before switching [44]. Here, we did not test for such a conformity bias based on the number of individuals using the same processing technique, but we rather tested for a conformity bias based on the frequency of technique observed. We thus tested whether individuals were i) more likely to use a technique frequently observed; ii) as likely to use a technique as it is observed or iii) less likely to use it. Such biases have been tested in great tits where the establishment of a foraging tradition has been found to rely on positive frequency-dependence; individuals adopting the most frequent local variant [30]. Barrett et al. [18] found no evidence for a conformity bias, but they found evidence of weak anti-conformity meaning that rare techniques attracted more attention in capuchins.

We found no evidence for kin- or sex-biased social learning. Previous studies reported that infant vervet monkeys ate the same food as their mother [29] and that females were preferred as models over males [26]. However, these studies tested for a single bias at a time while we tested here for several biases simultaneously, weighing each strategy against the other and compared to asocial learning which makes the analysis more powerful. Our results are furthermore consistent with a previous study testing for several model biases in a single analysis that did not identify such biases either [28]. We believe that testing for several biases in a single analysis allow to disentangle between potential confounding factors such as age, sex and rank and can highlight instances of equifinality where multiple social learning strategies produce the same signature in a population [45].

In the present study, the two innovators were an infant and a low-ranking adult male. In another open diffusion experiment [28], the alpha male and female were the two innovators in the same two groups when tested one year before. We believe that this difference can be linked to the aforementioned experiments’ methodological considerations. In Canteloup et al. study [28], groups were tested in a two-action box experiment for which individuals may get accustomed to boxes’ affordances despite the use of new design. Some studies suggested that juveniles and low rankers are often innovators [46, 47] although this depends on the type of behaviour [48], while other studies suggest neophilia and boldness have been found to better predict innovation [49]. We hypothesize that innovators’ identity also depends on task novelty. Monkeys being neophobic, an unknown food can be considered as a source of stress, which suggests that low rankers were the initiators and that juveniles were quicker to first succeed to extract peanuts because of their usual limited access to resources. When the task is not completely new, dominants are more expected to be the innovators.

We reported that, in the global model, social information influenced behaviour more heavily in adults than juveniles and in females than males, but these patterns were not consistent across all models. In the global and payoff-bias models, juveniles and females were most affected by payoff-based social information whereas adults and males were most affected by rank-based social information. Despite being in disagreement with what has been previously found in vervet monkeys [19], this suggests that juveniles and females would be more sensitive to behaviours’ payoffs while adults and males would be more sensitive to rank cues associated with a demonstrator. Overall, we found that adults weighed past experience more heavily than recent experience compared to juveniles, suggesting that memory strongly influences their behaviour as previously found in birds and capuchins [17, 18]. Vervet monkeys are used to extracting encased seeds from acacia pods like from pods of acacia nilotica (*Vachellia nilotica*). The habit of extracting seeds from these pods, although flatter and texturally different than unshelled peanuts, may influence the technique used by monkeys to open peanuts. However, personal experience prior to the experiment unlikely drives the diffusion of peanut opening techniques, as social learning models reliably predict its diffusion.

We show that payoff-biased social learning likely underlies the origin of the spread of novel food processing techniques in vervet monkeys, and that this content bias likely acts concurrently with rank-biased learning. EWA models consider exactly what behaviours and which demonstrators an individual uniquely observes both prior to their observing a successful solution and after they learn how to solve the task. This detailed time-varying modelling approach makes the analysis particularly powerful in uncovering the interplay between personal and social information use, which makes social learning adaptive and efficient [35]. Because it is very rare that the initial innovation event and its spread are recorded through observational studies, we believe that controlled experiments coupled with dynamic, theory-informed modelling are a useful approach to track the diffusion of new behaviours and to disentangle between different social learning strategies.

## Materials and Methods

### Study site and subject details

The study was conducted at the Inkawu Vervet Project (IVP) in a 12,000-hectare private game reserve: Mawana (28°00.327S, 031°12.348E) in KwaZulu Natal province, South Africa. The vegetation of the study site consisted in a savannah characterized by a mosaic of grasslands and clusters of trees of the typical savannah thornveld, bushveld and thicket patches. We studied two groups of wild vervet monkeys (*Chlorocebus pygerythrus*): ‘Noha’ (NH) and ‘Kubu’ (KB). NH was composed of 34 individuals (6 adult males; 9 adult females; 6 juvenile males; 7 juvenile females; 5 infant males; 1 infant female) and KB was composed of 19 individuals (1 adult male; 6 adult females; 3 juvenile males; 4 juvenile females; 3 infant males; 2 infant females; Table S1). Males were considered as adults once they dispersed, and females were considered as adults after they gave their first birth. Individuals that did not fulfil these criteria were considered as juveniles [28] and infants were aged less than one year old. IN EWA models, infants and juveniles were lumped in a single category “juveniles”. Each group had been habituated to the presence of human observers: since 2010 for NH and since 2013 for KB. All individuals were identifiable thanks to portrait photographs and specific individual body and face features (scars, colours, shape etc.).

This research adhered to the “Guidelines for the use of animals in research” of Association for Study of Animal Behaviour and was approved by the relevant local authority, Ezemvelo KZN Wildlife, South Africa.

### Hierarchy establishment

Agonistic interactions (aggressor behaviour: stare, chase, attack, hit, bite, take place; victim behaviour: retreat, flee, leave, avoid, jump aside) were collected from May 2018 to October 2018, aside from experiment days, on all the adults and juveniles of both groups via *ad libitum* sampling [50] and food competition tests (i.e. corn provided to the whole group from a plastic box). Data were collected by CC, MBC and different observers from the IVP team. Before beginning data collection, observers had to pass an inter-observer reliability test with 80% reliability for each data category between two observers. Data were collected on tablets (Vodacom Smart Tab 2) equipped with Pendragon version 8.

Individual hierarchical ranks were determined by the outcome of dyadic agonistic interactions recorded *ad libitum* and through food competition tests using Socprog software version 2.7 (50). Hierarchies in both groups were significantly linear (NH: h’ = 0.27; P < 0.0001; KB: h’ = 0.42; P < 0.0001) and ranks were assessed by I&SI method (52).

### Open diffusion experiment

The experimental apparatus consisted of provisioning the group with two transparent rectangular plastic boxes (34×14×12 cm) containing ∼ 2 kg unshelled peanuts in sufficient quantities to prevent a single individual from monopolizing the boxes. The monkeys were never provided with peanuts before the experiment and peanuts were not available in their environment. Thus, unshelled peanuts were a novel, nutritious food that required processing to be extracted from their shells before consumption.

Experiments took place at sunrise at monkeys’ sleeping sites during the dry, food-scarce winter to maximize their motivation for novel food. The two boxes of peanuts were offered to the monkeys, spaced apart by about 1 to 10 meters. CC led the experiment with the help of two to four field assistants. All monkeys were free to come to the boxes within the constraints of the social group dynamics. Experiments were video recorded using JVC cameras (EverioR Quad Proof GZ-R430BE) to which the experimenter said aloud the identities of the actor and of the attending neighbours for each manipulation event. A manipulation event was defined either as an attempt to extract a peanut from its shell (i.e. the individual acted on the peanut failing to fully open it and to get access to the food) or as a success (i.e. the individual succeeded to fully open a shell and to extract the peanut from the shell). A conspecific was considered as attending when it had its head or body oriented in an unobstructed line towards the demonstrator manipulating the peanut and was located within 0- 30m from the actor. Several individuals could thus be registered as attending to one or several demonstrators simultaneously.

The open diffusion experiments ran from May 2018 to August 2018 to maximise individuals’ likelihood of participating in the experiment. A total of 11 sessions of open diffusion experiments were run in NH and 10 in KB. The average duration of an experimental session was 46m:46s for NH and 42m:47s for KB.

### Video analysis

Video recordings were viewed and analysed by MBC with Media Player Classic Home Cinema software version 1.7.11. Twenty percent of the video were analysed by CC and inter-observer reliability was substantial (κ = 0.78). During video analysis in slow motion or frame by frame, the following variables were coded in an excel sheet: date, exact time of each manipulative event, actor identity, the technique used (crack with hand: ‘CH’; crack with mouth from the top of the peanut: ‘CMT’; crack with mouth from the side of the peanut: ‘CMS’; see Movies S1-S3) and the identity of attending individuals.

### Quantification and statistical analysis

Following Barrett et al. [17], we used a suite of hierarchical experience-weighted attraction (EWA) models to analyse data collected in the open-diffusion experiment. EWA models are time-series models that evaluate the joint influence of personal experience and social information on the probability of an individual displaying a behaviour [38] and are increasingly utilized in cross-taxa studies of cultural transmission [16-18, 27, 53].

This analytical approach has several strengths: It permits evaluation of multiple hypothesized learning strategies against each other and individual learning alone, utilizes a dynamic social learning network unique to each individual, and links individual variation in behaviour and cognition to population level-cultural dynamics. Working with time-series of behaviour unique to each individual is important as population-level signatures can often be misleading [54] or exhibit equifinality, particularly if individuals vary in experience, observation opportunities or the social learning strategies they employ [45]. The mathematical specification of our analytical approach also minimizes ambiguity of what types of social learning we are evaluating. This is important as verbal definitions are imprecise, and terminologies are differently interpreted in studies of social learning. Most importantly, this approach links theory to data. Instead of using a theoretically uninformed analytical approach to find results consistent with theory, we bypass quantitative proxies and translate theoretical models to statistical models. We fit a series of EWA models evaluating the following learning strategies:

1. Individual learning alone
2. Frequency-dependent learning (preference for behaviours that is based on their frequency in the population)
3. Female-biased learning (preference for the technique displayed by females in group i.e. matrilineal sex in vervets)
4. Matrilineal kin-biased learning (preference for the technique displayed by closely related individuals)
5. Compare means payoff-biased learning (preference for the most successful or efficient behaviour)
6. Rank-biased learning (preference for the technique displayed by high-ranking individual)
7. Sex-biased learning (preference for the behaviours of individuals that are of the same sex)
8. Global model that includes 1-7.

All social learning models (models 2-8) also include an individual learning component.

For each behavioural choice, social information used by an actor was the average value of each cue observed in a time window of 20 minutes prior to the observation (Table 1). As individuals access social information at different timescales, and this window choice was somewhat arbitrary, we also evaluated social info at 30, 10, and 5-minute timescales. These analyses yielded similar results and were robust to time windows.

We ran the EWA models using regularizing priors, which are sceptical of extreme effects and reduce the risk of overfitting, and a Cholesky decomposition for estimating varying effects. Models were fit using RStan version 2.19.3 [55]. Models were compared using widely applicable information criteria (WAIC), which can inform which model best predicts the observed data while penalizing models that underfit or overfit. Models with lower WAIC scores best predict the observed data.

### EWA Model Specification

EWA models have two parts: a set of expressions that specify how individuals accumulate experience and a second set of expressions that specify the probability of each option being chosen. Accumulated experience is represented by *attraction scores*, *A_ij,t_*, unique to each behaviour *i*, individual *j*, and time *t*. We update *A_ij,t_* with an observed payoff *π_ij,t_*:

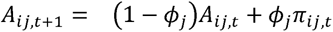

The parameter *ϕ*_j_ controls the importance of recent payoffs in influencing attraction scores. When *ϕ_j_* is high, more weight is given to recent experience over past experiences – memory has less of an influence on behavioural choice. This parameter is unique to an individual *j*, and we also estimate how it varies by age-class and sex.

Attraction scores are converted into probabilities of behavioural choice with a standard multinomial logistic choice rule:

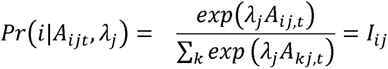

*λ_j_* controls sensitivity to differences in attraction scores on behavioural choice and is unique to an individual *j*. A very large *λ_j_*, means the option with the largest attraction score is nearly always selected. Choice is random with respect to attraction score when λ_j_ = 0. Individuals were assigned a payoff of zero, *π_ij,t_* = 0, if they failed to open a peanut. If they were successful *π_ij,t_* = 1.

Social learning may directly influence choice distinctly from individual learning. *S_ij_* = S(*i*|*Θ*_j_), is the probability an individual *j* chooses behaviour *i*, on the basis of a set of social cues and parameters *Θ_j_*. These social cues are traits associated with demonstrators (i.e. age, rank), or a behaviour (i.e. mean payoff), and each cue represents a hypothesized social learning strategy. Behavioural choice is a convex combination specified by:

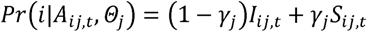

where is γ_j_ the weight assigned to social cues, and is bounded by 0 and 1.

Social cues are incorporated into *S_ij,t_*, by use of a multinomial probability expression with a log-linear component *B_ij,t_* that is an additive combination of cue frequencies. The probability of displaying each behaviour *i*, solely as a function of social cues, is:

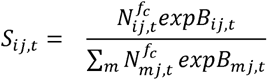

*N_ij,t_* are the observed frequencies of each technique *i* at time *t* by individual *j*. The exponentiated parameter *f_c_* controls the amount and type of frequency dependence. When *f_c_* = 1, social learning is unbiased by frequency and techniques influence choice in proportion to their occurrence (sometimes referred to as unbiased transmission). When *f*>1, social learning is positive frequency-dependent or conformist. When *f*<1, social learning is negative frequency-dependent, and a bias is shown towards rare behaviours.

Other social cues associate with individuals (i.e. rank, age, or relatedness) or behaviours (i.e payoffs), are incorporated via:

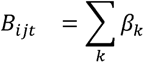

*B_ijt_* is the sum of the products of the influence parameters *β_k_* and the cue values *κ_k,ijt_*. Cues evaluated in these models are explained in detail in Table 1. The above specification is for the global model. Single social learning strategies are subset of this global model in which there is a single cue value or only frequency information is used.

### Other statistics

We used generalized linear models (GLMs) with quasi-Poisson error and log link function to test for the effect of group, normalized rank, sex and age on i) the latency of first success of shelling a peanut; ii) the number of peanut shelling successes; iii) the number of peanuts manipulation (attempts + successes); iv) the number of successes observed; v) the number of manipulations observed; vi) the number of times being observed when succeeding and vii) the number of times being observed when manipulating. We added the log of the time each individual had to process peanuts once they first succeeded as an offset (a standard statistical technique for converting a Poisson GLM for analysis of counts into a model for analysing counts per unit of time). All tests were performed with R Studio version 1.2.1335 using R version 3.6.1 [56].

## Acknowledgments

We thank Arend van Blerk and the whole IVP team for their help and support in the field. We are also particularly thankful to Loïc Brun, Adwait Deshpande, Manon Kerréveur-Lavaud, Maria Teresa Martinez Navarrete, Anaïs Rilly and Claudia Seminara for their assistance in data collection. We are grateful to the van der Walt family for their permission to conduct the study on their land. We warmly thank Matthias Wubs for writing us some R scripts to extract social data as matrices. We greatly thank Rachel Harrison for her comments on an initial version of our manuscript and the Editor and two anonymous reviewers for their comments that substantially improved our manuscript. This study has been funded by two postdoctoral fellowships from the Fyssen Foundation and the Fondation des Treilles granted to C.C. IVP was funded by the Swiss National Science Foundation (31003A_159587 and PP00P3_170624) and the Branco Weiss Fellowship – Society in Science granted to E.v.d.W.

## Author contributions

Data collection and video analysis: C.C. and M.B.C; statistical analysis: B.J.B (EWA) and C.C (GLMs); experimental design: C.C. and E.v.d.W; writing first draft: and B.J.B; reviews and editing: C.C., B.J.B. and E.v.d.W; funding: C.C. and E.v.d.W.

## Data availability

Reproducible model code and data can he found here: https://zenodo.org/record/4297318#.X8T5jC_pPUo

