## Supplemental Material for "Processing of novel food reveal payoff and rank-biased social learning in a wild primate"

CHARLOTTE CANTELOUP[1,2], MABIA BIFF CERA[1], BRENDAN J BARRETT[3,4,5]†, AND ERICA VAN DE WAAL[1,2]†

[1]Inkawu Vervet Project, Mawana Game Reserve, KwaZulu Natal, 3115, South Africa
[2]Department of Ecology and Evolution, University of Lausanne, 1015 Lausanne, Switzerland
[3]Max Planck Institute of Animal Behavior, Department for the Ecology of Animal Societies, Konstanz, Germany
[4]University of Konstanz, Department of Biology, Konstanz, Germany
[5]Max Planck Institute for Evolutionary Anthropology, Department of Human Behavior, Ecology, and Culture, Leipzig, Germany
†These authors contributed equally to this work

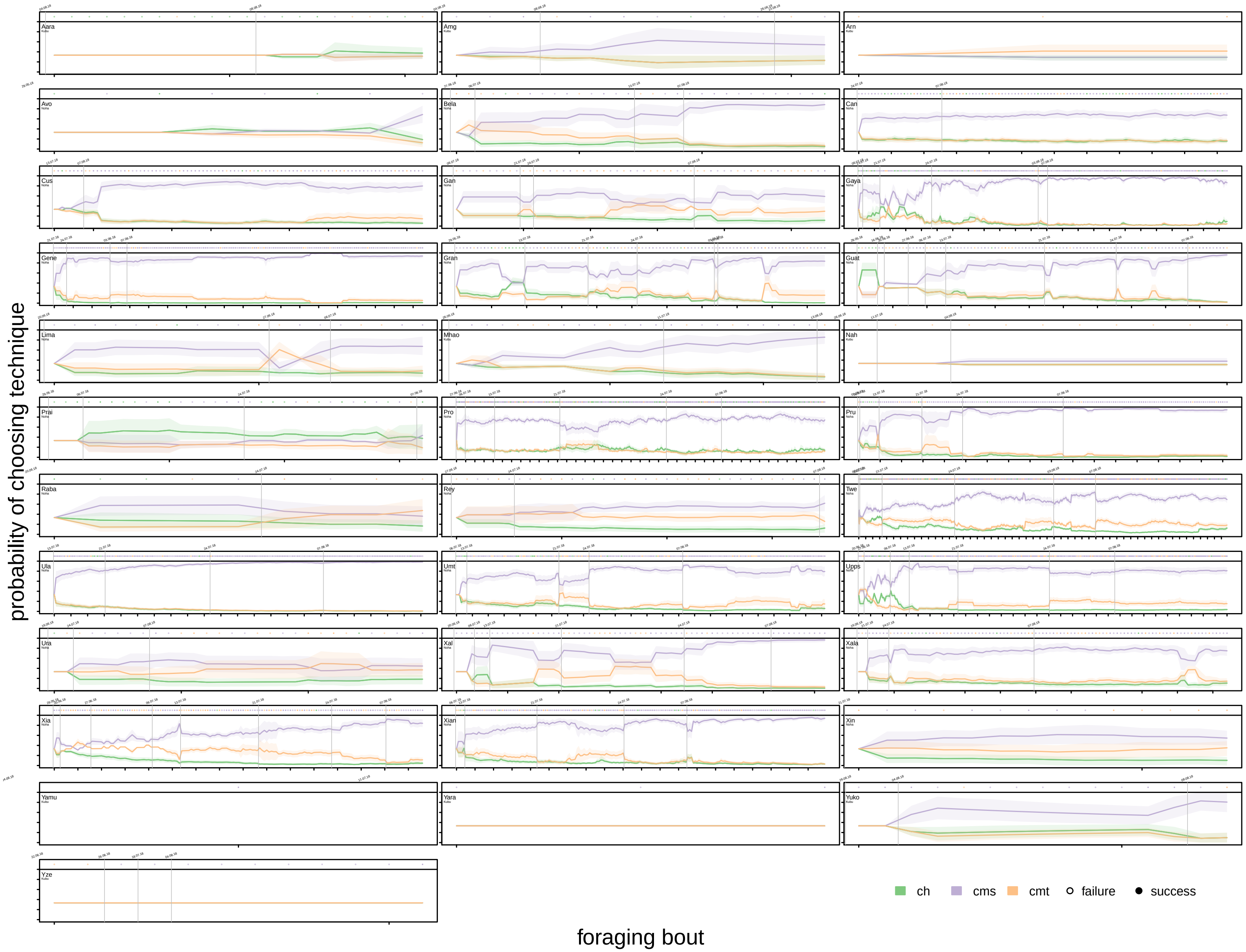


FIGURE S1. Model predictions of probability of choosing all 3 behavior at each foraging bout. Each panel is for a unique individual. Individual IDs are in top left corner, below that is individual group membership. Colors correspond to technique, line is posterior mean predictions corresponding to the probability of choosing a technique at each timestep, shaded area is 89% HPDI. Top row of each plot is raw data showing observed behavioral choice at each foraging bout. Filled circles correspond with successes, empty circles correspond with failures. Gray lines are drawn between experimental days.

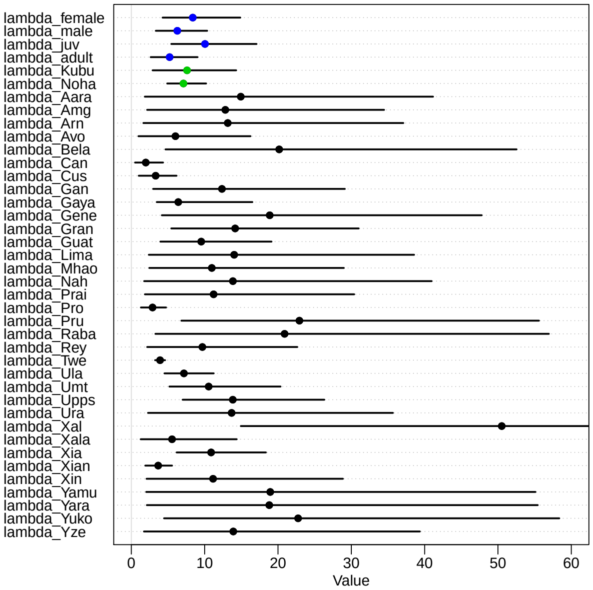


FIGURE S2. Dot plots of posterior predictions from the global model for all values of $\lambda$, the sensitivity to attraction scores. Points lie at posterior mean, line spans 89% HPDI. Blue points are main effects for each age and sex class, green points are varying effects of both groups, black points are individual-level varying effects. Values closer to zero indicate less sensitivity to differences in attraction scores.

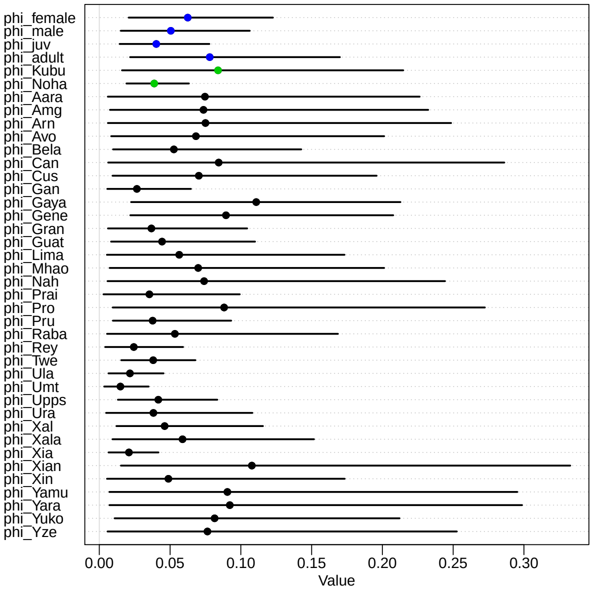


FIGURE S3. Dot plots of posterior predictions from the global model for all values of $\phi$, the weight given to recent experience. Points lie at posterior mean, line spans 89% HPDI. Blue points are main effects for each age and sex class, green points are varying effects of both groups, black points are individual-level varying effects. Lower values mean less weight given to recent experiences and a greater reliance on past memories.

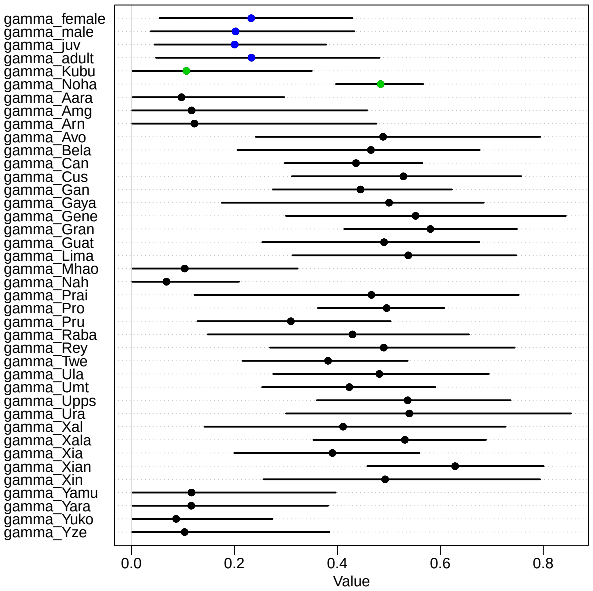


FIGURE S4. Dot plots of posterior predictions from the global model for all values of $\gamma$, the weight given to social information. Points lie at posterior mean, line spans 89% HPDI. Blue points are main effects for each age and sex class, green points are varying effects of both groups, black points are individual-level varying effects. Higher values indicate a grater reliance on social information than individual information.

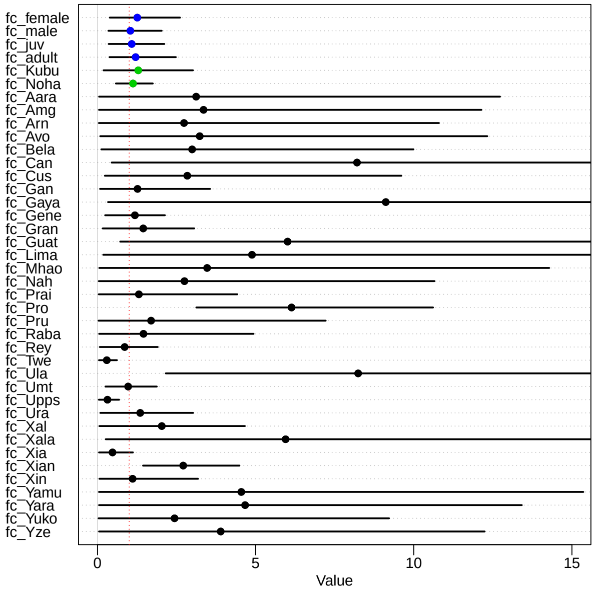


FIGURE S5. Dot plots of posterior predictions from the global model for all values of $f^c$, the strength of frequency dependence. Points lie at posterior mean, line spans 89% HPDI. Blue points are main effects for each age and sex class, green points are varying effects of both groups, black points are individual-level varying effects. Values $< 1$ are consistent with negative frequency-dependence, values of 1 indicate unbiased social learning, values $> 1$ indicate positive frequency dependence.

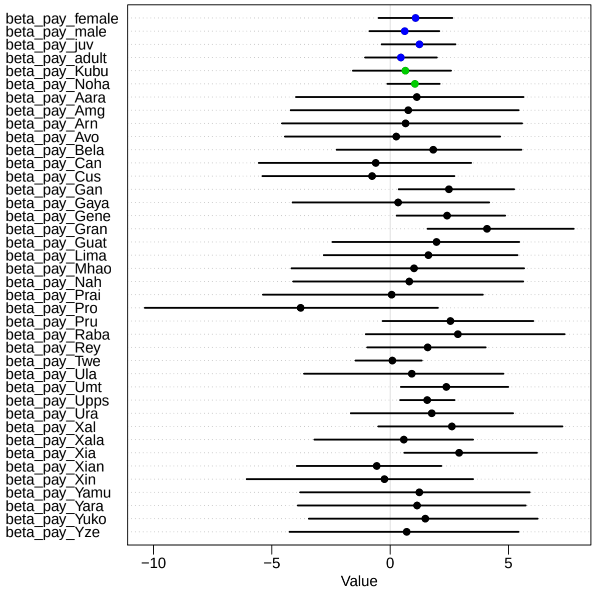


FIGURE S6. Dot plots of posterior predictions from the global model for all values of $\beta_{pay}$, the strength of payoff-bias. Points lie at posterior mean, line spans 89% HPDI. Blue points are main effects for each age and sex class, green points are varying effects of both groups, black points are individual-level varying effects.

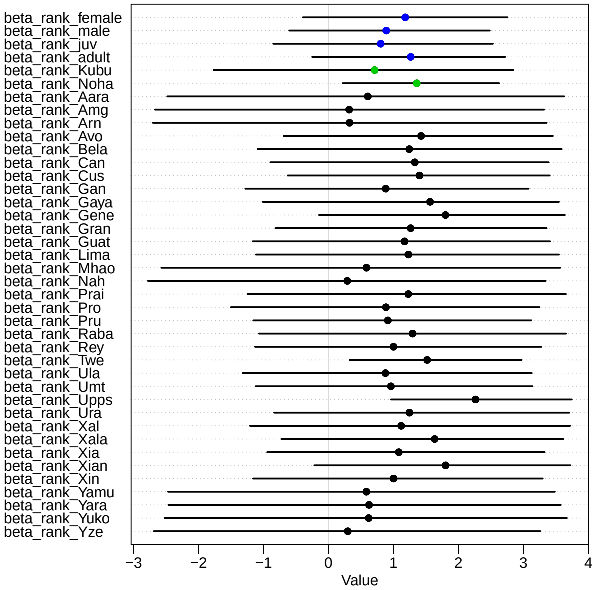


FIGURE S7. Dot plots of posterior predictions from the global model for all values of $\beta_{rank}$, the strength of payoff-bias. Points lie at posterior mean, line spans 89% HPDI. Blue points are main effects for each age and sex class, green points are varying effects of both groups, black points are individual-level varying effects.

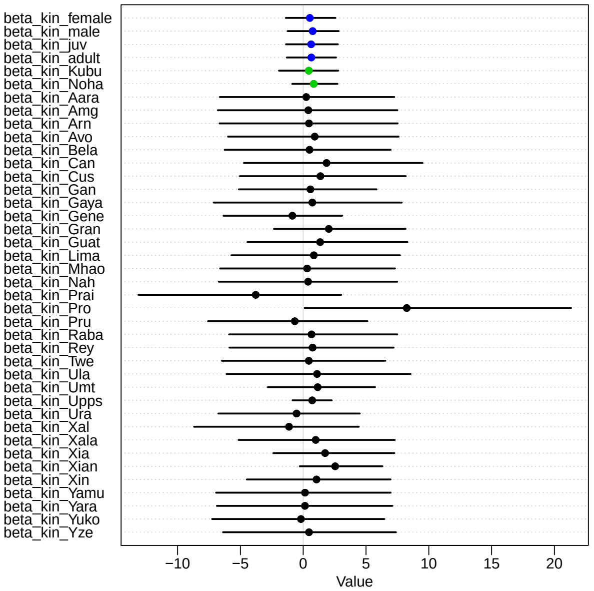


FIGURE S8. Dot plots of posterior predictions from the global model for all values of $\beta_{kin}$, the strength of payoff-bias. Points lie at posterior mean, line spans 89% HPDI. Blue points are main effects for each age and sex class, green points are varying effects of both groups, black points are individual-level varying effects.

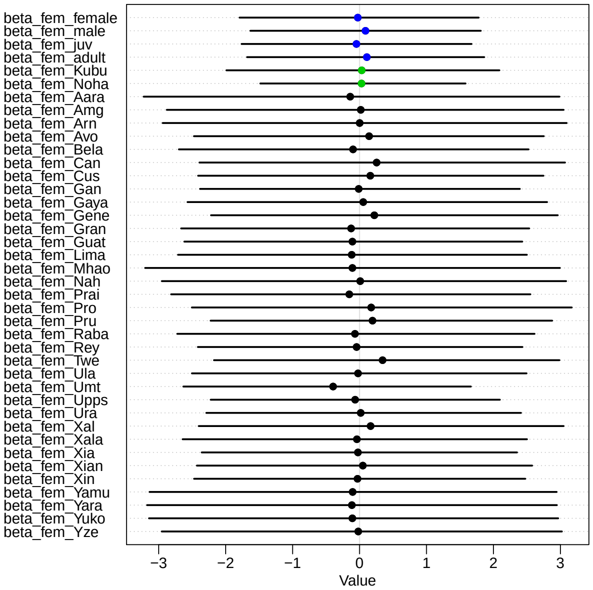


FIGURE S9. Dot plots of posterior predictions from the global model for all values of $\beta_{fem}$, the strength of payoff-bias. Points lie at posterior mean, line spans 89% HPDI. Blue points are main effects for each age and sex class, green points are varying effects of both groups, black points are individual-level varying effects.

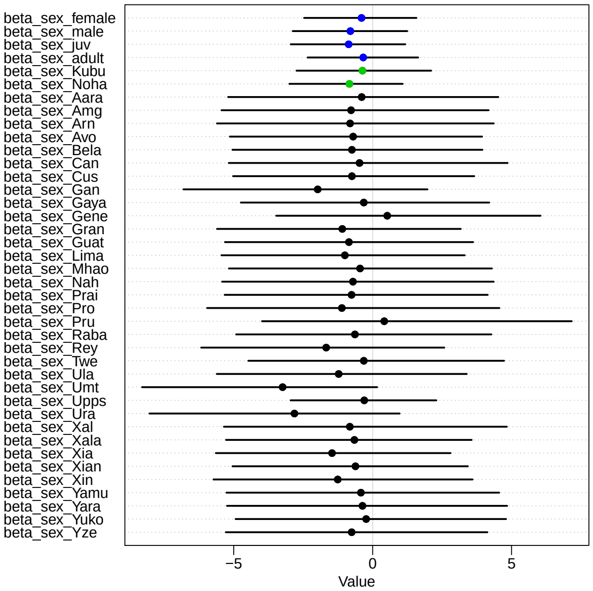


FIGURE S10. Dot plots of posterior predictions from the global model for all values of $\beta_{sex}$, the strength of payoff-bias. Points lie at posterior mean, line spans 89% HPDI. Blue points are main effects for each age and sex class, green points are varying effects of both groups, black points are individual-level varying effects.

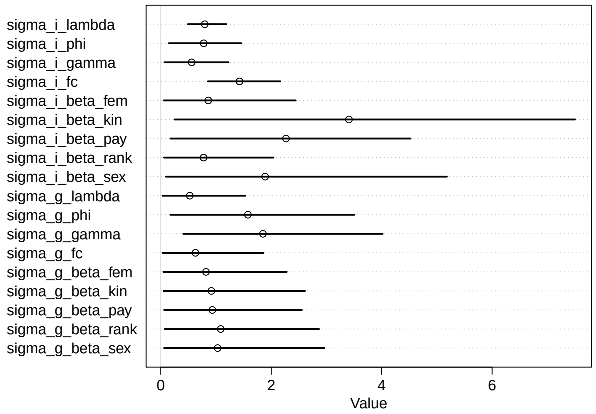


FIGURE S11. Dot plots of posterior predictions of variance, $\sigma$, for varying effects of parameters for the global model. Points lie at posterior mean, line spans 89% HPDI. $g$ is group level and $i$ is individual level variances.

| Group | Individual | Age | Sex | Rank | Std. rank | Latency first success | Order first success | First technique used | N success | N manipul. | N success observed | N manipul. observed | N success being observed | N manipul. being observed | Time available |
|---|---|---|---|---|---|---|---|---|---|---|---|---|---|---|---|
| Kubu | Aar | infant | male | NA | NA | 4737 | 1 | CMS | 16 | 44 | 0 | 3 | 29 | 81 | 20929 |
| Kubu | Aara | juvenile | female | 12 | 0.61 | 23616 | 9 | CH | 1 | 16 | 3 | 15 | 1 | 29 | 2050 |
| Kubu | Aare | adult | female | 7 | 0.33 | NA | NA | NA | NA | NA | 2 | 9 | NA | NA | NA |
| Kubu | Amg | infant | male | 14 | 0.72 | 19673 | 8 | CMS | 3 | 12 | 33 | 59 | 3 | 9 | 5993 |
| Kubu | Amur | adult | female | 8 | 0.39 | NA | NA | NA | NA | NA | 4 | 6 | NA | NA | NA |
| Kubu | Arn | juvenile | male | 6 | 0.28 | 6985 | 2 | CMT | 1 | 3 | 14 | 33 | 2 | 6 | 18681 |
| Kubu | Lif | adult | male | 1 | 0.00 | NA | NA | NA | NA | NA | 0 | 6 | NA | NA | NA |
| Kubu | Mal | juvenile | male | 17 | 0.89 | NA | NA | NA | NA | NA | 5 | 13 | NA | NA | NA |
| Kubu | Mara | adult | female | 15 | 0.78 | NA | NA | NA | NA | NA | NA | NA | NA | NA | NA |
| Kubu | Mhao | juvenile | female | 18 | 0.94 | 7011 | 3 | CMS | 16 | 25 | 10 | 14 | 22 | 22 | 18655 |
| Kubu | Mokc | infant | female | 16 | 0.83 | NA | NA | NA | NA | NA | 2 | 2 | NA | NA | NA |
| Kubu | Nah | juvenile | male | 10 | 0.50 | NA | NA | NA | NA | 11 | 4 | 8 | 0 | 14 | NA |
| Kubu | Ness | adult | female | 9 | 0.44 | NA | NA | NA | NA | NA | 1 | 9 | NA | NA | NA |
| Kubu | Yalu | adult | female | 2 | 0.06 | NA | NA | NA | NA | NA | 6 | 18 | NA | NA | NA |
| Kubu | Yamu | juvenile | female | 3 | 0.11 | 18650 | 7 | CMS | 1 | 1 | 2 | 8 | 1 | 1 | 7016 |
| Kubu | Yara | infant | female | 13 | 0.67 | 13953 | 5 | CMS | 1 | 3 | 1 | 2 | 2 | 4 | 11713 |
| Kubu | Yeni | adult | female | 4 | 0.17 | NA | NA | NA | NA | NA | 0 | 1 | NA | NA | NA |
| Kubu | Yuko | juvenile | female | 5 | 0.22 | 7413 | 4 | CMS | 5 | 15 | 4 | 20 | 4 | 7 | 18253 |
| Kubu | Yze | infant | male | 11 | 0.56 | 18570 | 6 | CMS | 1 | 12 | 1 | 5 | 1 | 19 | 7096 |
| Noha | Avo | adult | male | 25 | 0.73 | 122 | 1 | CMS | 2 | 6 | 5 | 11 | 12 | 24 | 30744 |
| Noha | Bela | juvenile | female | 24 | 0.70 | 11707 | 11 | CMT | 13 | 31 | 95 | 159 | 16 | 51 | 19159 |
| Noha | Can | adult | male | 13 | 0.36 | 24270 | 24 | CMS | 73 | 114 | 290 | 464 | 188 | 299 | 6596 |
| Noha | Cus | adult | male | 16 | 0.45 | 17876 | 19 | CMS | 68 | 94 | 107 | 172 | 94 | 132 | 12990 |
| Noha | Gan | infant | male | 18 | 0.52 | 23137 | 23 | CMS | 13 | 56 | 153 | 222 | 36 | 138 | 7729 |
| Noha | Gaya | adult | female | 4 | 0.09 | 13996 | 15 | CMS | 141 | 236 | 218 | 348 | 588 | 881 | 16870 |
| Noha | Gene | adult | female | 3 | 0.06 | 20861 | 20 | CMS | 144 | 196 | 310 | 476 | 239 | 322 | 10005 |
| Noha | Gran | juvenile | female | 1 | 0.00 | 159 | 2 | CMS | 51 | 108 | 264 | 402 | 172 | 369 | 30707 |
| Noha | Guat | juvenile | female | 6 | 0.15 | 6187 | 5 | CMS | 37 | 109 | 175 | 277 | 110 | 306 | 24679 |
| Noha | Lima | juvenile | female | 26 | 0.76 | 9354 | 6 | CMS | 13 | 26 | 46 | 96 | 11 | 39 | 21512 |
| Noha | Prai | juvenile | female | 12 | 0.33 | 2158 | 4 | CH | 14 | 33 | 44 | 72 | 17 | 44 | 28708 |
| Noha | Pret | adult | female | 11 | 0.30 | NA | NA | NA | NA | NA | 26 | 50 | NA | NA | NA |
| Noha | Pro | juvenile | male | 19 | 0.55 | 11538 | 10 | CMS | 225 | 403 | 485 | 769 | 333 | 617 | 19328 |
| Noha | Pru | juvenile | male | 23 | 0.67 | 13985 | 14 | CMS | 87 | 174 | 435 | 709 | 147 | 274 | 16881 |
| Noha | Pye | infant | male | 31 | 0.91 | NA | NA | NA | NA | NA | 16 | 41 | NA | NA | NA |
| Noha | Raba | juvenile | female | 32 | 0.94 | 9760 | 8 | CMS | 4 | 9 | 28 | 59 | 1 | 2 | 21106 |
| Noha | Renn | adult | female | 27 | 0.79 | NA | NA | NA | NA | NA | 4 | 8 | NA | NA | NA |
| Noha | Reva | adult | female | 34 | 1.00 | NA | NA | NA | NA | NA | 4 | 4 | NA | NA | NA |
| Noha | Rey | juvenile | male | 30 | 0.88 | 11791 | 12 | CMT | 21 | 36 | 63 | 110 | 47 | 70 | 19075 |
| Noha | Rioj | infant | female | 33 | 0.97 | NA | NA | NA | NA | NA | 5 | 16 | NA | NA | NA |
| Noha | Roma | adult | female | 28 | 0.82 | NA | NA | NA | NA | NA | 5 | 6 | NA | NA | NA |
| Noha | Rosl | juvenile | female | 29 | 0.85 | NA | NA | NA | NA | NA | 5 | 9 | NA | NA | NA |
| Noha | Twe | adult | male | 2 | 0.03 | 16388 | 17 | CMS | 374 | 431 | 112 | 198 | 1489 | 1996 | 14478 |
| Noha | Ula | juvenile | male | 10 | 0.27 | 16954 | 18 | CMS | 156 | 222 | 434 | 722 | 452 | 630 | 13912 |
| Noha | Umt | juvenile | male | 14 | 0.39 | 13938 | 13 | CMS | 139 | 233 | 566 | 911 | 290 | 491 | 16928 |
| Noha | Upps | adult | female | 7 | 0.18 | 9806 | 9 | CMT | 200 | 312 | 401 | 658 | 569 | 998 | 21060 |
| Noha | Ura | infant | male | 21 | 0.61 | 26886 | 25 | CMS | 6 | 30 | 476 | 715 | 24 | 115 | 3980 |
| Noha | Wol | adult | male | 5 | 0.12 | NA | NA | NA | NA | NA | 8 | 11 | NA | NA | NA |
| Noha | Xal | infant | male | 20 | 0.58 | 9573 | 7 | CMS | 16 | 73 | 142 | 233 | 44 | 168 | 21293 |
| Noha | Xala | adult | female | 8 | 0.21 | 21090 | 22 | CMS | 54 | 105 | 217 | 335 | 217 | 402 | 9776 |
| Noha | Xia | juvenile | male | 17 | 0.48 | 173 | 3 | CMS | 79 | 157 | 163 | 260 | 207 | 419 | 30693 |
| Noha | Xian | adult | female | 9 | 0.24 | 14823 | 16 | CH | 138 | 200 | 222 | 386 | 485 | 638 | 16043 |
| Noha | Xin | infant | male | 22 | 0.64 | 21056 | 21 | CMT | 8 | 14 | 218 | 357 | 30 | 52 | 9810 |
| Noha | Yan | adult | male | 15 | 0.42 | NA | NA | NA | NA | NA | 27 | 64 | NA | NA | NA |

TABLE S1. Composition of the two study groups Noha and Kubu. Individual level variables (individual, age, sex, rank, normalized rank); latency of first peanut opening success (in seconds); order of first peanut opening success; first technique used; number of successes; number of manipulations (successes + attempts); number of successes observed; number of manipulations observed; number of times being observed when succeeding; number of times being observed when manipulating; time available over the whole experiment after the first success (in seconds). NA indicates individuals did not manipulate or succeed to open peanuts.
